# γ-Amino Carboxylic Acid Modification Enhances the Efficacy of PNAs Targeting miR-221-3p in A549 Cells

**DOI:** 10.1101/2025.06.15.659802

**Authors:** Youngsim Yoon, Na-rae Joo, Junghyun Min, Daeyoon Bae, Seohee Lee, Youngjun Choi

## Abstract

Peptide nucleic acids (PNAs) are versatile tools for diagnostic and therapeutic applications, including gene regulation and miRNA targeting. However, their therapeutic potential is often limited by challenges such as biological efficacy. To address this, PNAs offer a key advantage over other nucleic acids—their ease of modification, which allows for enhanced properties. In this study, we introduced a novel *γ*-amino carboxylic acid (*γ*-ACA) modification to PNAs targeting miR-221-3p, a key miRNA implicated in various pathological processes. The modified PNAs showed significantly improved binding affinity to their targets and more efficient inhibition of miR-221-3p expression compared to unmodified PNAs in A549 cells, leading to effective regulation of downstream gene and protein expression. These results highlight the potential of *γ*-modified PNAs as a platform for developing miRNA-targeted therapeutics.

## Introduction

PNAs have emerged as powerful tools for a diverse range of applications, including gene silencing, diagnostics, and therapeutic interventions^1–3^. Their unique structure, characterized by a backbone made of peptide-like bonds instead of the conventional sugar-phosphate backbone found in natural nucleic acids, confers distinct advantages such as enhanced binding affinity, stability, and specificity^1,2,4^. Unlike traditional DNA or RNA, PNAs possess a neutral charge, eliminating electrostatic repulsion during hybridization and allowing for more flexible binding conditions^2,5^. These characteristics make PNAs especially suitable for targeting genetic mutations and dysregulated gene expression in diseases like cancer, underlining their strong potential for therapeutic use^6–8^.

Although PNAs show great potential, their practical application, especially in therapy, is limited by several challenges. One major challenge is their poor aqueous solubility and low cellular uptake efficiency, primarily attributable to their neutral charge and limited interaction with lipid-rich cell membranes^7–9^. Moreover, achieving therapeutic effects with PNAs often requires high doses due to their dose-dependent activity, rapid systemic clearance, poor tissue distribution, and limited cellular permeability^10–12^. This high dosing requirement complicates clinical translation and poses challenges regarding potential side effects from repeated administrations. Additionally, PNAs are prone to aggregation in solution, further restricting their use in biological systems^13^. To address these challenges, structural modifications—such as introducing charged groups into the PNA backbone—have been explored to improve solubility, stability, and cellular uptake^14^.

*γ*-modified PNAs (*γ*-PNAs) introduce a stereocenter at the *γ* carbon of the PNA backbone^15^. This modification enhances PNA solubility and cellular uptake, while simultaneously increasing their binding affinity for DNA or RNA targets^16–18^. The right-handed helical conformation restricts molecular flexibility, leading to more stable and selective binding interactions^7^,^16^,^18^. For example, *γ*-guanidinyl-modified PNAs (*γ*-GPNAs) enhance binding affinity and Hoogsteen-face selectivity for miRNA targets. Additionally, they prevent undesired ternary complex formation. This modification also facilitates internalization into living cancer cells, suggesting its potential for therapeutic applications^18^. Similarly, *γ*-GPNAs exhibit improved hybridization with complementary DNA and RNA due to their structural preorganization and the presence of guanidine groups, which significantly enhance cellular uptake^16^. Furthermore, *γ*-PNAs with miniPEG side chains improve aqueous solubility while retaining strong nucleic acid binding and low cytotoxicity, rendering them effective in targeting double-stranded DNA via Watson–Crick base pairing^19^. Such modifications have been successfully employed to silence oncogenic microRNAs, including miR-210, leading to significant tumor growth inhibition *in vivo* when delivered via PLGA nanoparticles^17^. These developments highlight the potential of *γ*-PNAs for applications in various fields, including gene therapy, diagnostics, and nanotechnology.

In line with, we introduce a novel *γ*-amino carboxylic acid (*γ*-ACA) modification to enhance PNA performance further. This modification is designed to target miR-221-3p, a key microRNA implicated in cancer and other diseases^20–23^. It demonstrates significantly stronger binding affinity and more effective suppression of miR-221-3p expression in A549 cells compared to unmodified PNAs. Notably, this suppression is accompanied by upregulation of p27, a tumor suppressor protein, and its encoding gene, *CDKN1B*, indicating a broader regulatory impact on downstream gene expression^24^,^25^. These findings establish *γ*-ACA-modified PNAs (*γ*-ACA-PNAs) as a promising therapeutic platform for targeted gene regulation. Our work marks a significant step toward more effective miRNA-targeted therapies and advances the field of precision medicine in cancer treatment.

## Results

### Synthesis of *γ*-ACA PNAs

The chemical structures of unmodified PNA and *γ*-ACA-PNA are compared in Fig. 1, the synthetic route for *γ*-ACA-PNA is outlined in Fig. 2, where the *γ*-ACA monomer was synthesized through a multi-step process and incorporated into the PNA backbone via solid-phase synthesis.

**Figure 1.**
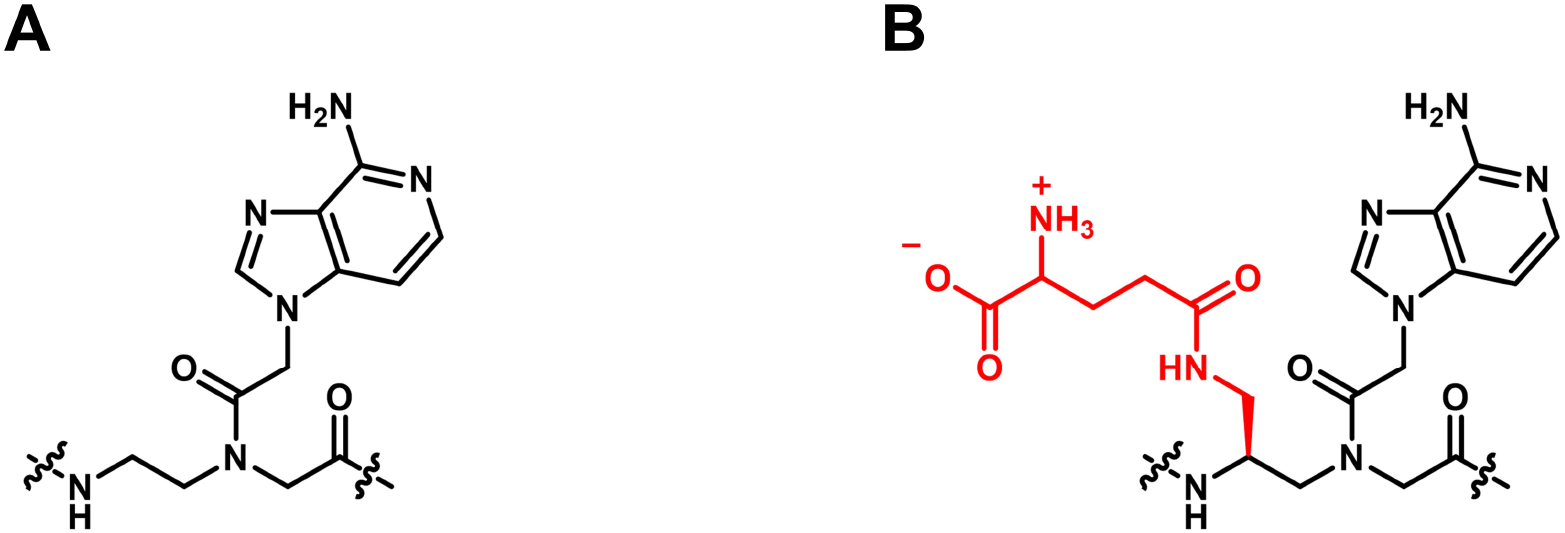
Chemical structure of (A) aeg PNA-A; (B) *γ*-(S-amonocarboxylic acid) aeg PNA-A

**Figure 2.**
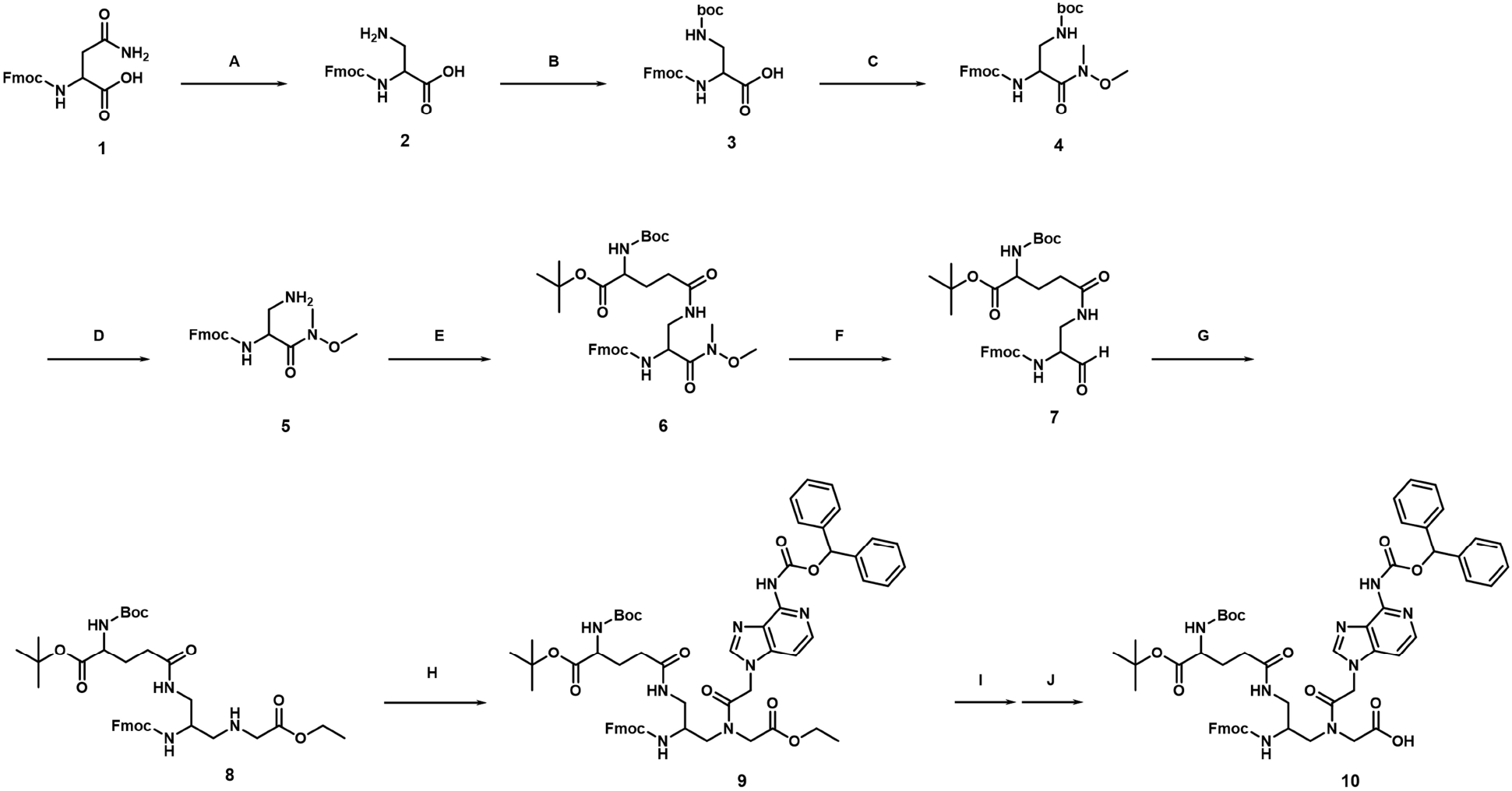
(A) Iodosobenzene diacetate, EA/MeCN/H2O, 4hr at RT, 70%; (B) (Boc)2O, NaHCO3, THF/H2O, 3hr at RT, 95%; (C) N,O-dimethyl hydroxylamine.HCl, HBTU, DIEA, DMF, 2hr at RT, 85%; (D) TFA/MC, 1.5hr at RT (Crude); (E) Boc-Glu-OtBu, HBTU, DIEA, DMF, 4hr at RT, 92%; (F) 2M LiAlH4 in THF, THF(HPLC), 3hr at -55oC, 97%; (G) Gly-OEt.HCl, AcOH, NaBH3CN, MeOH, 2hr at RT, 46%; (H) A(Bhoc)-aecetic acid, DhbtOH, EDC.HCl, DMF, 2hr at RT, 71%; (I) 1N LiOH in H2O/THF, 30min at 5oC; (J) Fmoc-OSu, 10% Na2CO3, 2hr at RT, 98%

In the initial stage, Fmoc-Asn-OH was used as the starting material, which was oxidized using (Diacetoxyiodo)benzene to obtain the amine derivative. This was followed by Boc protection and O-acylation reactions, leading to the synthesis of compounds 3 and 4. Subsequently, Weinrab amide compound 5 was synthesized using N,O-dimethylhydroxylamine and tetramethyluronium hexafluorophosphate (HBTU), and Boc-Glu-OtBu was introduced through an amide bond formation (amide coupling) reaction.

Next, a partial reduction reaction using 2M LiAlH_4_ in tetrahydrofuran (THF) was performed to convert the intermediate into an aldehyde (-CHO). This was followed by a reductive amination reaction using glycine ethyl ester hydrochloride (Gly-OEt·HCl) and NaBH_3_CN, leading to the successful synthesis of the key intermediate (compound 8). This then incorporates both amine (-NH_2_) and carboxylic acid (-COOH) functional groups at the *γ*-position.

From this key intermediate, various base acetic acid derivatives were utilized in amide bond formation reactions, enabling the synthesis of PNA monomer derivatives with diverse basic substituents. In the final stage, hydrolysis and N-terminal Fmoc protection using Fmoc-Osu were performed, leading to the successful synthesis of various final PNA monomers. Following the successful synthesis of these *γ*-ACA-PNA monomers, they were subsequently used for the synthesis of PNA oligomers, including those with unmodified monomers.

### Characteristics of *γ*-ACA Modified PNAs

To explore the impact of *γ*-ACA modification on target binding affinity, melting temperature (Tm) measurements were conducted using new PNA oligomers designed to target miR-221-3p (Table 1)^26^. The *γ*-ACA modification was applied to PNAs of varying lengths (18mer, 15mer, and 12mer) to assess its universal applicability. The results revealed that *γ*-ACA modification significantly increased Tm values for both DNA and RNA targets, with the 18mer *γ*-ACA PNA showing an increase of 7.4 ^*°*^C for DNA and 7.5 ^*°*^C for RNA compared to the unmodified PNA. Similar trends were observed for the 15mer and the 12mer PNAs, indicating a consistent stabilizing effect of *γ*-ACA modification across different PNA lengths. These trends were consistently observed in both UV spectrophotometry and fluorescence-based qPCR analysis, reinforcing the stabilizing effect of *γ*-ACA modification on PNA-target duplexes. Collectively, these findings highlight the potential of *γ*-ACA modification to improve PNA performance for therapeutic applications.

**Table 1.**
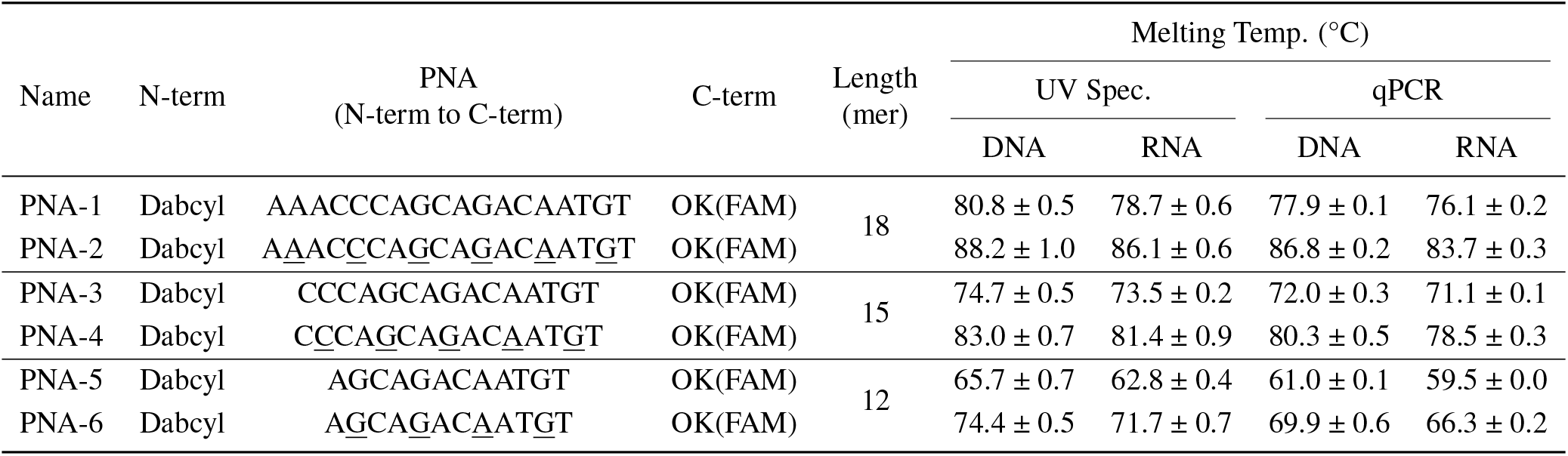
Sequences and Melting temperature (Tm) of PNA oligomers. The *γ*-ACA modified sequences are highlighted with an underline.

To assess whether *γ*-ACA modification influences cellular uptake, flow cytometry analysis was performed on A549 cells treated with unmodified PNA (PNA-1) and *γ*-ACA-PNA (PNA-2). The results showed that *γ*-ACA modification alone had almost no effect on cellular uptake, with fluorescence levels similar to those of unmodified PNA (Fig. 3A). To enhance intracellular delivery, the Tat-modified peptide (RRRQRRKKRR), a type of cell-penetrating peptide (CPP), was conjugated to *γ*-ACA-PNA (PNA-7), resulting in a substantial increase in cellular uptake. PNA-7 treatment led to complete internalization in all A549 cells, confirming that Tat-modified peptide conjugation plays a crucial role in enhancing cellular delivery of *γ*-ACA-PNA (Fig. 3B).

**Figure 3.**
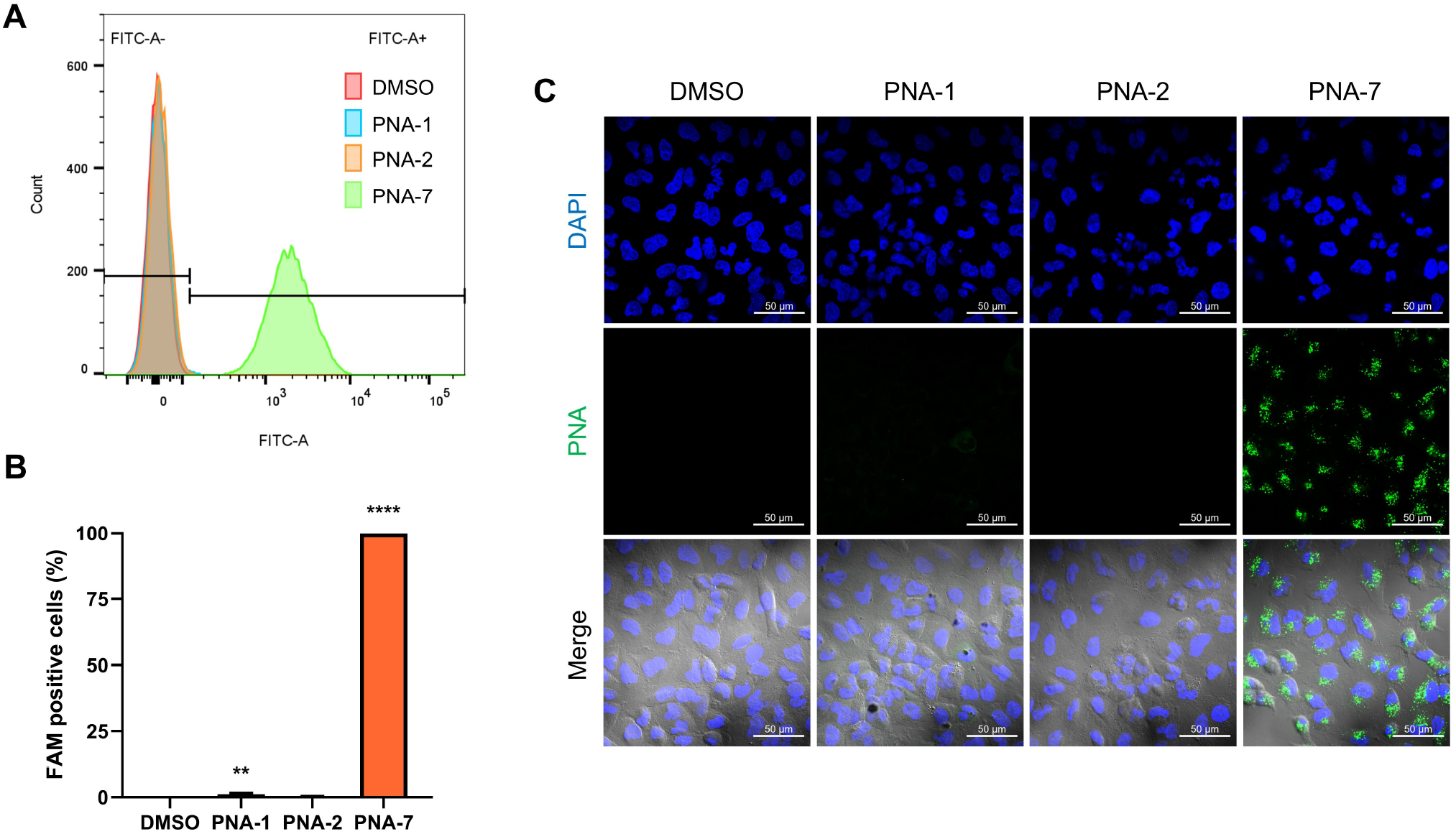
Evaluation of Cellular Uptake of *γ*-ACA Modified PNAs in A549 Cells (A) A representative histogram showing the intracellular delivery efficiency of PNA-1, PNA-2, and PNA-7 (5 *µM*) for 24 hr and (B) quantification of FAM-positive A549 cells analyzed by flow cytometer. 10,000 events were recorded for each sample. (C) Confocal microscopy images of A549 cells treated with PNA-1, PNA-2, and PNA-7 (5 *µM*) for 24 hr. Scale bar, 50 *µm*. Results were reported as mean ± SD (n=3). Statistical analysis : **P < 0.01, ****P < 0.0001 (One-way ANOVA followed by Tukeyś test)

### Evaluation of miRNA Inhibition by *γ*-ACA Modification

To further investigate the functional effect of *γ*-ACA modification, miRNA inhibition assays were conducted using PNA-8 to PNA-13, which were selected to eliminate unnecessary components such as fluorophores and quenchers, retaining only essential elements for precise inhibition analysis, and a scramble PNA (PNA-SC) as a negative control (Table 2). A549 cells were treated with 5*µM* of each PNA for 24 hours, and miR-221-3p levels were quantified.

**Table 2.**
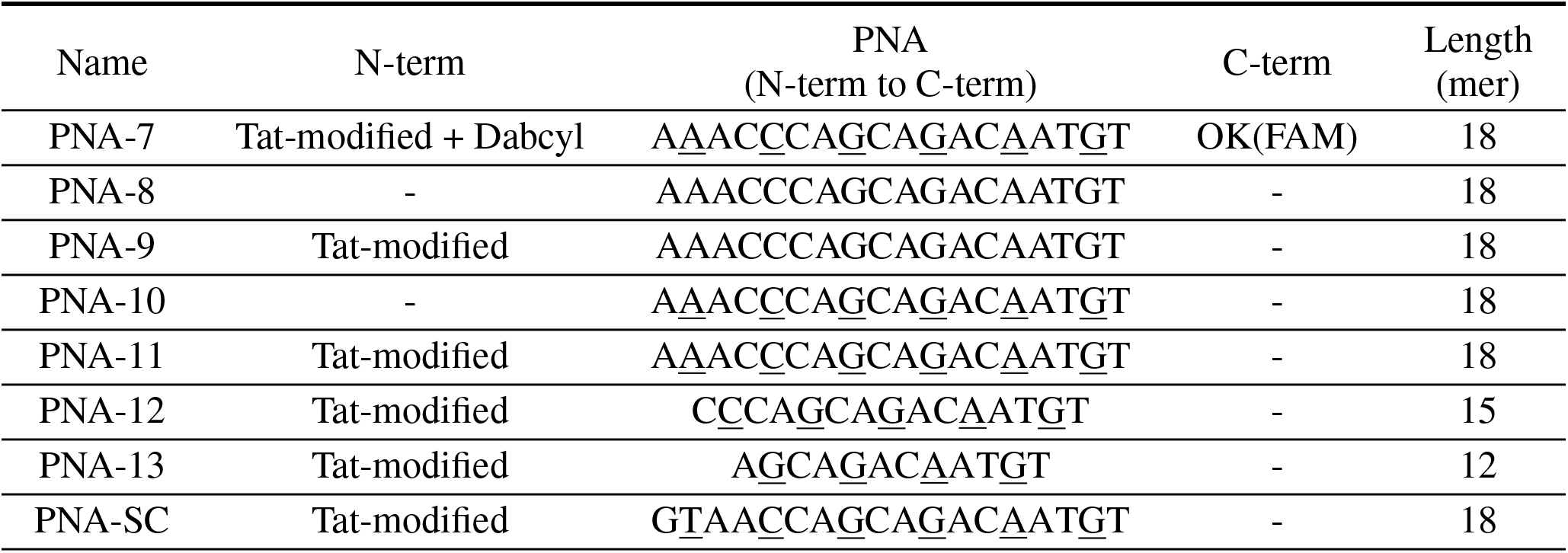
PNA oligomers with modified sequences targeting miR-221-3p. The sequences with *γ*-ACA modifications are highlighted with an underline.

PNA-8 (unmodified PNA) achieved a modest reduction in miR-221-3p levels and exhibited the lowest inhibition efficiency among the tested PNAs (Fig. 4A). PNA-9, which incorporated the Tat-modified peptide conjugation into the unmodified PNA, showed a slight improvement in inhibition, indicating that CPP conjugation enhances PNA efficacy. Notably, PNA-10 (*γ*-ACA-modified PNA) exhibited a more substantial increase in miRNA inhibition compared to PNA-8 and PNA-9, suggesting that *γ*-ACA modification contributes more significantly to enhancing PNA activity than the increase in cellular uptake mediated by CPP conjugation. Among the tested PNAs, PNA-11, which incorporated both *γ*-ACA and the Tat-modified peptide, showed the most pronounced inhibitory effect, highlighting the synergistic impact of these modifications. In contrast, the scramble PNA (PNA-SC) demonstrated no effect, confirming the specificity of the *γ*-ACA-PNAs for miR-221-3p.

**Figure 4.**
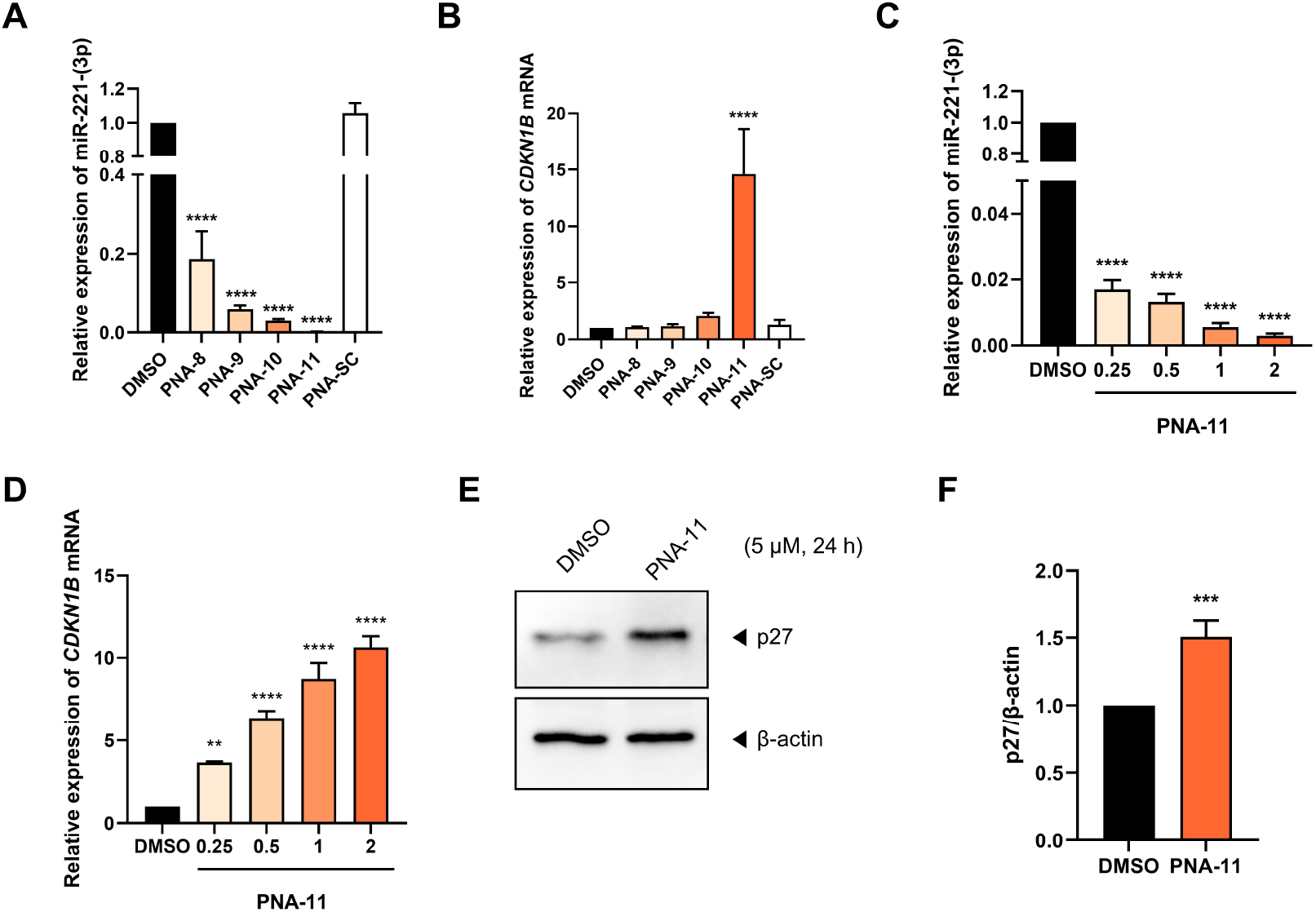
miR-221-3p Inhibition and Downstream Effects of *γ*-ACA Modified PNAs (A) Fold changes of miR-221-3p and (B) *CDKN1B* mRNA after treatment with PNA-8, PNA-9, PNA-10, PNA-11, and PNA-SC (5 *µM*) for 24 hr. (C) Fold changes of miR-221-3p and (D) *CDKN1B* gene expression of PNA-11 with different concentration (0.25, 0.5, 1, and 2 *µM*) for 24 hr. (E) Representative western blot images showing the expression of p27 in A549 cells treated with 5 *µM* of DMSO or PNA-11 for 24 hr. (F) Quantification of p27 expression levels normalized to *β* -actin. Results were presented as mean ± SD (n=3). Statistical analysis: **P < 0.01, ***P < 0.001, ****P < 0.0001 vs. DMSO treated group (One-way ANOVA followed by Tukey’s test or unpaired Student’s t-test)

The downstream effects of miR-221-3p inhibition were also assessed by measuring the expression of *CDKN1B*, a key gene regulated by miR-221-3p. Consistent with the miRNA inhibition data, *CDKN1B* mRNA expression was dramatically upregulated in the cells treated with PNA-11, which was not observed with the other PNAs, compared to the untreated control (Fig. 4B). Importantly, this upregulation was dose-dependent, with PNA-11 showing miRNA inhibition and *CDKN1B* induction even at lower concentrations (0.25-2*µM*), as illustrated in Fig. 4C and D. Western blot analysis further confirmed these findings, showing a marked increase in the expression of p27 protein, encoded by *CDKN1B*, in response to PNA-11 treatment (Fig. 4E and F). Collectively, these results underscore the ability of *γ*-ACA modification to enhance both miRNA inhibition and the activation of downstream targets, positioning it as a powerful tool for therapeutic applications.

### Influence of PNA Length on miRNA Inhibition

To determine the effect of PNA length on miR-221-3p inhibition, A549 cells were treated with *γ*-ACA-modified PNAs of varying lengths (PNA-11: 18mer, PNA-12: 15mer, PNA-13: 12mer) at a concentration of 5*µM* for 24 hours. The PNAs were designed by retaining the C-terminal region, which targets the 5’ end of miR-221-3p, while sequentially truncating the N-terminal region, as the 5’ end contains the seed sequence, crucial for miRNA target recognition and binding^27,28^. The results indicated a clear correlation between PNA length and inhibitory efficiency. Specifically, the 18mer PNA (PNA-11) exhibited the highest miRNA inhibition rate, while the 15mer (PNA-12) showed a significant inhibition, and the 12mer (PNA-13) exhibited the weakest inhibition (Fig. 5A). These findings suggest that longer PNAs may achieve stronger target binding and better inhibitory effects due to their increased hybridization potential.

**Figure 5.**
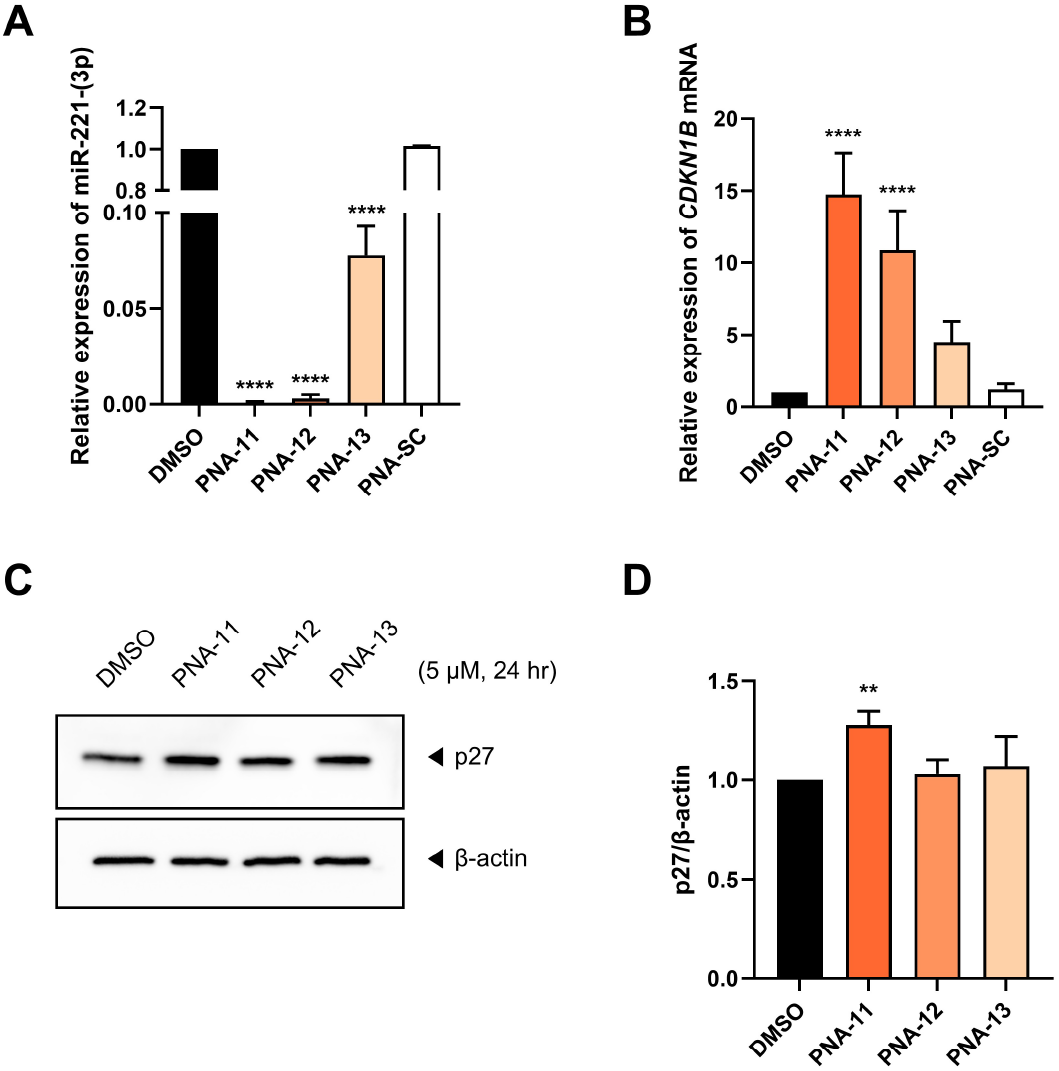
Impact of PNA Length on miR-221-3p Inhibition and Downstream Gene Expression (A) Fold change of miR-221-3p expression after PNA-11, PNA-12, PNA-13, and PNA-SC (5 *µM*) treatment for 24 hr. miR-16-5p was used as control. (B) Relative expression of CDKN1B after PNA-11, PNA-12, PNA-13, and PNA-SC (5 *µM*) treatment for 24 hr. GAPDH was used as a control. (C) Representative western blot images of p27 expression level treated with DMSO, PNA-11, PNA-12, or PNA-13 (5 *µM*) for 24 hr, and (D) Quantification of p27 protein levels based on western blot band intensities. Results were reported as mean ± SD (n=3). Statistical analysis: **P < 0.01, ****P < 0.0001 vs. DMSO treated group (One-way ANOVA followed by Tukey’s test or unpaired Student’s t-test)

Consistent with the miRNA inhibition data, *CDKN1B* expression was most significantly upregulated in cells treated with PNA-11, followed by a substantial increase in PNA-12-treated cells, and the least increase was observed in cells treated with PNA-13 (Fig. 5B). Western blot analysis further demonstrated that p27 protein levels were significantly increased only in response to PNA-11 treatment (Fig. 5C and D). This suggests that shorter PNAs may be insufficient to effectively inhibit miR-221-3p and elicit significant biological responses, emphasizing the critical role of PNA length in achieving robust gene regulation.

### PNA-11 Induces Ferroptosis via miR-221-3p Inhibition

Given the strong inhibitory effect of PNA-11 on miR-221-3p and the associated upregulation of *CDKN1B*, we next sought to determine whether this molecular modulation leads to cell death. Annexin V and propidium iodide staining revealed significant increases in both apoptotic and necrotic cell populations following PNA-11 treatment, suggesting the involvement of multiple cell death pathways (Fig. 6A, 6B, and 6C). The increase in apoptotic cells is consistent with the upregulation of *CDKN1B*, a well-known cell cycle inhibitor and pro-apoptotic regulator.

**Figure 6.**
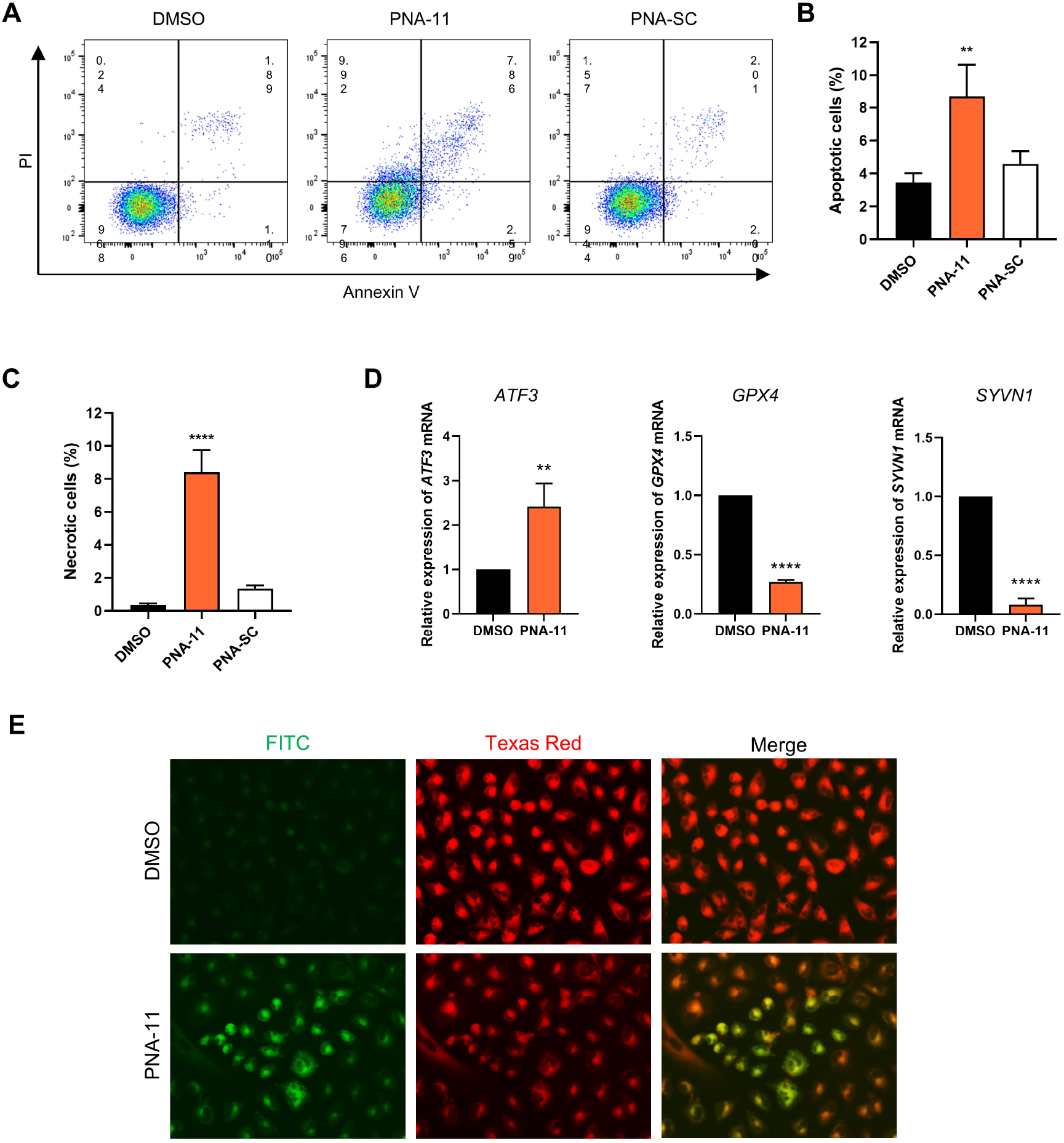
(A) Flow cytometry analysis of apoptotic cells treated with PNA-11 and PNA-SC (5 *µM*) treatment for 72 hr. Quantification of (B) apoptotic cells and (C) necrotic cells based on flow cytometry data. (D) Relative expression levels of *ATF3, GPX4*, and *SYVN1* mRNA after PNA-11 and PNA-SC (5 *µM*) treatment for 24 hr. GAPDH was used as a control. (E) Representative images of C-11-BODIPY staining after treatment with DMSO and PNA-11 (5 *µM*) for 48 hr. Results were reported as mean ± SD (n=3). Statistical analysis: **P < 0.01, ****P < 0.0001 vs. DMSO treated group (One-way ANOVA followed by Tukey’s test or unpaired Student’s t-test)

In parallel, the elevated necrotic cell population prompted us to explore the possible involvement of ferroptosis, a distinct form of regulated necrosis characterized by iron-dependent lipid peroxidation. Previous studies have reported the involvement of miR-221-3p in the regulation of ferroptosis^29^. To investigate whether ferroptosis was responsible for the observed necrotic phenotype, we first examined the expression of ferroptosis-related genes. The results showed a marked increase in *ATF3*, a validated target of miR-221-3p and a known promoter of ferroptosis, whereas the expression levels of *GPX4* and *SYVN1*—genes known to suppress ferroptosis—were significantly reduced following PNA-11 treatment (Fig. 6D). To further support these findings, C11-BODIPY staining was conducted. A significant increase in lipid ROS accumulation was observed in PNA-11-treated cells, indicative of ferroptosis induction (Fig. 6E).

Taken together, these findings demonstrate that the *γ*-ACA-modified PNA not only efficiently suppresses miR-221-3p and upregulates its downstream target *CDKN1B*, but also induces both apoptosis and ferroptosis in A549 cells, highlighting its strong potential as a multifunctional therapeutic agent.

## Discussion

The role of PNAs in modulating miRNA expression has been an area of increasing interest, as part of ongoing efforts to enhance gene regulation strategies. In this study, we demonstrated that a newly designed PNA oligomer with *γ*-amino carboxylic acid (*γ*-ACA) modification effectively inhibits miR-221-3p expression in A549 cells. This modification significantly improved target binding affinity, leading to enhanced suppression of miR-221-3p and subsequent upregulation of *CDKN1B* expression. Notably, despite the relatively poor cellular uptake of *γ*-ACA-modified PNA, our data demonstrated that it exhibited superior target inhibition compared to regular PNA conjugated with the Tat-modified peptide. Previous studies have suggested that conjugation with cell-penetrating peptides can enhance the miRNA inhibition efficacy of PNAs^30^; however, our results indicate that the *γ*-ACA modification alone is sufficient to achieve greater target suppression. These findings suggest that optimizing binding affinity may be a more effective strategy than improving cellular uptake for enhancing PNA efficacy.

Our data indicate that the observed increase in *CDKN1B* expression following miR-221-3p inhibition supports the hypothesis that the modified PNA effectively disrupts miRNA-mediated suppression. Since *CDKN1B* is a critical regulator of cell cycle progression, and its coding protein, p27, plays a key role in cellular homeostasis. These results suggest potential applications for *γ*-ACA-modified PNAs in therapeutic strategies targeting miRNA dysregulation. Furthermore, compared to conventional PNAs, which often require additional conjugation strategies to enhance efficacy, our findings suggest that strategic modifications aimed at strengthening PNA-miRNA interactions could be a viable alternative approach for optimizing therapeutic outcomes.

*γ*-PNAs have been investigated for their potential applications in various biological contexts. A chemically modified *γ*-PNA encapsulated in PLGA nanoparticles was developed to inhibit miR-210, a key regulator in cervical cancer and other solid tumors^17^. Treatment with this *γ*-PNA significantly reduced tumor growth and improved tumor morphology in a HeLa xenograft model. Similarly, another study employed tail clamp *γ*-PNA targeting miR-155 in lymphoma cells and a xenograft mouse model, demonstrating a 90–95% reduction in miR-155 levels, increased apoptosis, and significant tumor growth suppression^31^. Beyond miRNA targeting, *γ*-PNAs have also been applied to genomic DNA targeting, showing efficacy in silencing oncogenes such as c-Myc in multiple cancer models^32^. These findings highlight the broad applicability of *γ*-PNA modifications beyond miRNA inhibition, suggesting their potential use in gene regulation and cancer therapy.

In addition to cancer therapy, *γ*-PNAs have shown promise in the field of gene editing. *γ*-PNAs with polyethylene glycol substitutions have been used to enhance DNA binding, facilitating efficient gene editing in hematopoietic stem cells both *ex vivo* and *in vivo*, with clinically relevant levels of correction observed in models of *β* -thalassemia^33^. Furthermore, *γ*-PNAs with hydroxymethyl-*γ*-substituents have demonstrated superior gene-editing efficiency compared to conventional PNAs, further emphasizing their potential for therapeutic applications^34^.

Our study uniquely demonstrates that *γ*-ACA modification selectively strengthens binding affinity while maintaining biological efficacy. This enhancement is likely attributable to structural stabilization and improved molecular interactions with the target miRNA. Modifications at the *γ*-position of PNA have been shown to induce helical preorganization, transforming randomly folded structures into stable helices, which may contribute to increased binding affinity.

Specifically, the *γ*-ACA modification may promote additional hydrogen bonding or alter the conformational rigidity of the PNA backbone, facilitating stronger hybridization with its target. Structural studies on PNA-dsRNA triplexes have identified hydrogen bonding interactions between the PNA amide hydrogen and RNA phosphate oxygen, suggesting that *γ*-ACA modifications could similarly enhance hydrogen bonding interactions with target miRNA^7^,^9^,^15^. These structural adaptations may collectively contribute to the enhanced hybridization efficiency and miRNA inhibition observed in our study, underscoring the significance of *γ*-ACA modification in optimizing PNA function.

Despite these promising findings, certain limitations should be acknowledged. First, our study was conducted in an *in vitro* model using A549 cells, and *in vivo* validation is necessary to assess the therapeutic potential of the modified PNA comprehensively. Furthermore, advanced delivery systems must be explored to optimize biodistribution and intracellular delivery. Given that *γ*-ACA modification primarily enhances binding affinity, complementary delivery strategies—such as nanoparticle-based carriers or lipid conjugation—could further improve therapeutic performance.

The exact molecular mechanisms by which *γ*-ACA modification enhances PNA properties remain to be fully elucidated. Structural analysis and biophysical characterization of binding dynamics using techniques such as isothermal titration calorimetry (ITC) or molecular dynamics (MD) simulations are required to better understand these mechanisms. Additionally, it is crucial to evaluate the potential off-target effects of *γ*-ACA-modified PNAs to ensure their specificity and minimize unintended interactions with non-target sequences.

In conclusion, our study provides novel insights into the potential of *γ*-ACA-modified PNAs as effective miRNA inhibitors. By demonstrating that improved target binding affinity directly contributes to enhanced gene regulation, our findings contribute to the advancement of PNA-based approaches for gene modulation. Further investigations into the structural and functional properties of this modification will be essential for optimizing PNA design and expanding its therapeutic applications.

## Materials and Methods

### Chemicals for PNA synthesis

(6-Benzhydryloxycarbonylamino-purin-9-yl)-acetic acid (A(Bhoc)-acetic acid), (2-Benzhydryloxycarbonylamino-6-oxo-1,6-dihydro-purin-9-yl)-acetic acid (G(Bhoc)-acetic acid), (4-N-(Benzhydryloxycarbonyl)cytosine)-1-acetic acid (C(Bhoc)-acetic acid), Thymine-1-acetic acid were synthesized by HLB Panagene (Daejeon, Republic of Korea).

All chemical products were purchased through Sigma-Aldrich (St. Louis, MO, USA). Those chemical products includes 2M Lithium aluminium hydride in tetrahydrofuran (LiAlH in THF), Sodium cyanoborohydride (NaBH_3_CN), N-*α*-Fmoc-L-asparagine (Fmoc-Asn-OH), (Diacetoxyiodo)benzene, Di-tert-butyl dicarbonate((Boc)_2_O), N,N,N^*′*^, N^*′*^ - Tetramethyl-O-(1H-benzotriazol-1-yl)uronium hexafluorophosphate,O-(Benzotriazol-1-yl)-N,N,N^*′*^, N^*′*^-tetramethyluronium hexafluorophosphate (HBTU), N,O-dimethylhydroxylamine·HCl,N-(3-Dimethylaminopropyl)-N^*′*^-ethylcarbodiimide hydrochloride (EDC·HCl), Boc-L-glutamic acid 1-tert-butyl ester (Boc-Glu-OtBu), 9-Fluorenylmethyl-succinimidyl carbonate (Fmoc-Osu), 3,4-Dihydro-3-hydroxy-4-oxo-1,2,3-benzotriazine (HODHBt), Trifluoroacetic acid (TFA), Acetonitrile (ACN), Ethyl acetate (EA), Dichloromethane (DCM), N,N-dimethylformamide (DMF), Diethyl ether (Et_2_O), tetrahydrofuran (THF), Methanol, Sodium bicarbonate (NaHCO_3_), Sodium carbonate(Na_2_CO_3_), Magnesium sulfate(MgSO_4_), 1N Hydrochloric (1N HCl), N,N-diisopropylethylamine (DIEA), Acetic acid, Lithium hydroxide (LiOH) and Glycine ethyl ester hydrochloride.

### Synthesis of PNA oligomers

The PNA oligomers used in this study were synthesized on a solid support (Rink amide AM Resin) using standard Fmoc chemistry with an automatic synthesizer (HLB Panagene, Daejeon, Republic of Korea), incorporating both unmodified Fmoc PNA monomers and *γ*-ACA-PNA monomers^35–37^. The synthesized oligomers were cleaved from the resin using a cocktail solution of m-cresol and trifluoroacetic acid (TFA) (1:4), followed by precipitation with ether, purification, and analysis by reverse-phase high-performance liquid chromatography (RP-HPLC) and matrix-assisted laser desorption/ionization time-of-flight (MALDI-TOF) mass spectrometry.

### Cell culture and PNA transfection

A549 human lung adenocarcinoma cells were obtained from the Korean Cell Line Bank (KCLB, Seoul, South Korea) and cultured in RPMI-1640 medium (Cytiva, DE, USA) supplemented with 10% fetal bovine serum (FBS, Cytiva) and 1% penicillin/streptomycin (10,000 U*/*mL penicillin and 10,000 U*/*mL streptomycin; Cytiva) in a humidified CO_2_ incubator at 37 ^*°*^C. For PNA transfection, A549 cells were seeded in a 6-well plate and cultured overnight. The culture medium was replaced with a fresh medium containing PNA at the indicated concentration. After transfection, the cells were harvested and used for subsequent experiments.

### Thermal melting experiments

The melting temperatures were measured using UV spectrophotometry and quantitative polymerase chain reaction (qPCR). For UV spectrophotometry, PNA (4.5 *µM*) and its complementary DNA (4.5 *µM*; IDT, Coralville, IA, USA) or RNA (4.5 *µM*; IDT) were mixed at a 1:1 ratio in 1 ×Tris-EDTA buffer under PNA:DNA and PNA:RNA conditions, and absorbance was measured at 260 nm with a Cary 3500 Multicell UV-Vis Spectrophotometer (Agilent, Santa Clara, CA, USA). The temperature was decreased from 95 ^*°*^C to 45 ^*°*^C at a rate of 2 ^*°*^C/min, and measurements were performed in triplicate. For qPCR, PNA (0.4 *µM*) and its complementary DNA (0.4 *µM*, IDT) or RNA (0.4 *µM*, IDT) were mixed at a 1:1 ratio and analyzed with the QuantStudio 5 real-time PCR System (Applied Biosystems, Foster City, CA, USA). The temperature was increased from 40 ^*°*^C to 99.9 ^*°*^C°C at a rate of 0.5 ^*°*^C every 5 seconds, with triplicate measurements to collect data.

### Transfection efficiency

A549 cells were subcultured in either 6-well plates or 4-well glass slides and incubated for 24 hours before PNA treatment. After 24 hours of incubation with PNA, the inoculation medium was removed, and cells were washed three times with either sterile Phosphate-buffered saline (PBS) or Hanks’ balanced salt solution (HBSS). For flow cytometry analysis, cells from the 6-well plates were detached using trypsin, washed with PBS, and then suspended in cold PBS containing 10% fetal bovine serum. The samples were analyzed using a BD FACSCanto II flow cytometer (BD Biosciences, San Jose, CA, USA.), with quantification performed using FlowJo software v10 (BD Biosciences). For confocal microscopy, cells on glass slides were stained with NucBlue™ Live ReadyProbes™ Reagent (Thermo Fisher Scientific, Waltham, MA, USA) and CellMask™ Deep Red Actin Tracking Stain (Invitrogen, Carlsbad, CA, US), incubated at 37 ^*°*^C with 5% CO_2_ for 15 minutes, washed three times with HBSS media, fixed with 4% paraformaldehyde (PFA), and then observed using LSM 700 confocal microscopy (Carl Zeiss, Oberkochen, Germany).

### RNA extraction

Modified PNA was transfected into A549 cells, and after 24 hours, the cells were harvested using TRIzol™ Reagent (Invitrogen) following the manufacturer’s instructions. The extracted RNA was dissolved in nuclease-free water and quantified using a NanoDrop One Microvolume UV-VIs spectrophotometer (Thermo Fisher Scientific) for subsequent experiments.

### Reverse transcription and Quantitative real-time PCR

To quantify the expression levels of miR-221-3p and mRNA trasncripts in A549 cells, cDNA synthesis was first performed using the miRNA 1st-Strand cDNA Synthesis Kit (Agilent) for miR-221-3p analysis, with the Universal reverse primer (Agilent) and the RevertAid First Strand cDNA Synthesis Kit (Thermo Fisher Scientific) for mRNA transcripts analysis, both following the manufacturer’s protocols with the C1000 Touch Thermal Cycler (Bio-Rad, Hercules, CA, USA). Quantitative polymerase chain reaction (qPCR) was then conducted using 20 *×* EvaGreen™ (BIOFACT, Daejeon, South Korea) on the QuantStudio 5 real-time PCR System (Applied Biosystems). The qPCR was performed in triplicate to confirm the results, with miR-16-5p used as the internal control miRNA for miR-221-3p quantification and Glyceraldehyde 3-phosphate dehydrogenase (GAPDH) used as the reference control for normalization of gene expression.

The primers used for miRNA qPCR were as follows: The forward primer for miR-221-3p was TAGCAGCACG-TAAATATTGGCG, and the forward primer for the internal control miR-16-5p was AGCTACATTGTCTGCTGGGTTTC; both were used in combination with the Universal reverse primer provided in the Agilent miRNA quantification system.

For mRNA quantification, the primer pairs were as follows: *CDKN1B* forward TAATTGGGGCTCCGGCTAACT and reverse TGCAGGTCGCTTCCTTATTCC; *ATF3* forward GCTGGAAAGTGTGAATGCTG and reverse TTCTGAGCCCGGA-CAATACA; *GPX4* forward GTAACCAGTTCGGGAAGCAG and reverse TGTCGATGAGGAACTGTGGA; *SYVN1* forward CCAACATCTCCTGGCTCTTTCAC and reverse GTCAGGATGCTGTGATAGGCGT; *GAPDH* forward CTGGGCTACACT-GAGCACC and reverse AAGTGGTCGTTGAGGGCAATG.

### Western blot analysis

A549 cells were washed with ice-cold PBS and lysed in radioimmunoprecipitation assay (RIPA) buffer supplemented with a protease inhibitor (Thermo Fisher Scientific). Total protein lysates (30*µg*) were loaded onto a 4–20% Mini-PROTEAN TGX precast gel (Bio-Rad) and electrophoresed at 100 V for 90 minutes. The separated proteins were transferred to polyvinylidene difluoride (PVDF) membranes (Bio-Rad) at 100 V for 90 minutes. The membranes were blocked with a blocking reagent for 1 hour at room temperature and incubated overnight at 4°C with anti-*β* -actin (mouse monoclonal, 1:5,000; Sigma-Aldrich, St. Louis, MO, USA) or anti-p27 Kip1 (rabbit monoclonal, 1:2,000; Cell Signaling Technology, Danvers, MA, USA) primary antibodies. After washing with Tris-buffered saline with 0.1% Tween 20 (TBST), the membranes were incubated with horseradish peroxidase (HRP)-conjugated secondary antibodies (1:2,000; Abcam, Cambridge, UK) for 1 hour at room temperature. Protein bands were imaged using SuperSignal West Pico PLUS Chemiluminescent Substrate (Thermo Fisher Scientific) and ChemiDoc Imaging System (Bio-Rad).

### Analysis of Apoptosis and Necrosis

Apoptotic cell analysis was performed using the Dead Cell Apoptosis Kit with Annexin V for Flow Cytometry (Thermo Fisher Scientific), according to the manufacturer’s instructions. Briefly, cells were trypsinized, washed with PBS, and resuspended in Annexin V binding buffer. Cells were then incubated with Annexin V and propidium iodide (PI) at room temperature for 15 minutes. After incubation, Annexin V binding buffer was added to each sample, and the cells were analysed using a BD FACS Canto II flow cytometer (BD Biosciences). Quantification of apoptotic and necrotic cells was carried out using FlowJo software version 10 (BD Biosciences).

### Detection of Ferroptosis by BODIPY™ 581/591 C11

A549 cells were seeded in a 6-well plate and incubated for 24 hours. After 24 hours, PNA was added to each well at a final concentration of 5 *µM*, and the cells were incubated for an additional 48 hours. Following the 48-hour treatment, the culture medium was removed, and the cells were labeled with BODIPY™ 581/591 C11 (Invitrogen) diluted in complete RPMI medium to a final concentration of 2 *µM*. The cells were incubated at 37 ^*°*^C in a 5% CO_2_ incubator for 30 minutes. After incubation, the cells were washed three times with sterile PBS and observed under a fluorescence microscope.

### Statistical analysis

Comparisons between two groups were analyzed using unpaired Student’s t-test, while comparisons among three or more groups were performed using one-way ANOVA. A P-value of < 0.05 was considered statistically significant. Data were presented as group mean ± SD. Statistical analyses were conducted using Prism version 10 (GraphPad Software, San Diego, CA, USA). Statistical significance was defined as follows: *, P < 0.05; **, P < 0.01; ***, P < 0.001; ****, P < 0.0001

## Acknowledgments

We gratefully acknowledge Haejin Jung, Principal Technical Officer at the Research Solution Center (RSC), Institute for Basic Science (IBS), for performing the flow cytometry analysis.

## Author contributions statement

Y.Y. and N.J. conceived and performed the experiments and contributed to manuscript preparation. J.M. designed and synthesized the PNA monomer. D.B. and S.L. synthesized and purified the PNA monomer and oligomer. Y.C. contributed to the experimental design, data analysis, and manuscript preparation. All authors reviewed and approved the final manuscript.

## Additional information

